# Shoelaces: an interactive tool for ribosome profiling processing and visualization

**DOI:** 10.1101/279497

**Authors:** Åsmund Birkeland, Katarzyna Chyżyńska, Eivind Valen

## Abstract

The emergence of ribosome profiling to map actively translating ribosomes has laid the foundation for a diverse range of studies on translational regulation. The data obtained with different variations of this assay is typically manually processed, which has created a need for tools that would streamline and standardize processing steps.

We present Shoelaces, a toolkit for ribosome profiling experiments automating read selection and filtering to obtain genuine translating footprints. Based on periodicity, favoring enrichment over the coding regions, it determines the read lengths corresponding to bona fide ribosome protected fragments. The specific codon under translation (P-site) is determined by automatic offset calculations resulting in sub-codon resolution. Shoelaces provides both a user-friendly graphical interface for interactive visualisation in a genome browser-like fashion and a command line interface for integration into automated pipelines. We process 79 libraries and show that studies typically discard excessive amounts of data in their manual analysis pipelines.

Shoelaces streamlines ribosome profiling analysis offering automation of the processing, a range of interactive visualization features and export of the data into standard formats. Shoelaces stores all processing steps performed in an XML file that can be used by other groups to exactly reproduce the processing of a given study. We therefore anticipate that Shoelaces can aid researchers by automating what is typically performed manually and contribute to the overall reproducibility of studies. The tool is freely distributed as a Python package, with additional instructions and demo datasets available at https://bitbucket.org/valenlab/shoelaces

## Background

Ribosome profiling provides the first opportunity to monitor the behavior of translating ribosomes on a transcriptome-wide scale. Since its development [1], the technique has been widely adopted and inspired a diverse range of studies on translational regulation. While the assay itself has been partially standardized, the processing of data has not. A significant bottleneck is that of reproducibility and interpretation. In particular, most studies relies on manual selection of read lengths and manual P-site determination. The choices made are highly variable between studies and make it challenging to compare results in the literature.

The consistent processing of such data necessitates that two major challenges are met: (1) separating signal from noise, i.e. distinguishing footprints of translating ribosomes from reads originating from other processes and (2) determining the specific codon being translated by the ribosome which the read fragment originates from (a P-site offset). While some software tools have been developed for analyzing ribosome profiling data (for an overview see [2]), few address these challenges directly. Instead, tools typically rely on manual selection of read lengths and offsets or perform selection as part of an integrated pipeline with no data export options [3, 4, 5].

Here, we introduce Shoelaces, a software tool for processing ribosome profiling data. Shoelaces addresses the processing challenges by (1) utilizing a property of phasing, a strong 3-nucleotide periodicity of the reads stemming from coding regions [1, 6, 7] to filter genuine translating footprints and (2) calibrating P-site offsets based on metagene profiles over start or stop codons, stratified by footprint length [1, 8]. Shoelaces automatically selects these lengths and offsets, as well as offers batch-mode for processing multiple libraries in bulk.

The tool can be run in two modes: either using a graphical or command line interface. The graphical interface is accessible to users of all levels and guides the user through each processing step, allows for interactive adjustments and offers a range of extra visualization features on both gene/transcript or global level. The command line interface offers the same functionality as the graphical interface, without the interactivity, and can be easily integrated into automated processing pipelines.

## Implementation

Shoelaces is implemented in Python3 and designed to run on Linux and MacOS operating systems. It relies on OpenGL for rendering graphics and PyQt5 for cross-platform graphical user interface. GUI is composed of a set of windows that user can easily rearrange by drag-and-drop to customize layout. The plots are interactive making the processing easily adjustable to specific needs. While primarily designed for the visualization features, Shoelaces can be also run in command line, making it easy to incorporate into processing pipelines. Shoelaces operates on common genomic formats (BAM, GTF, BED, wiggle), and stores settings in XML files, for maximum ease of use and reproducibility of analyses.

## Results and discussion

### Data processing workflow

The workflow of Shoelaces is shown in Fig. 1. Shoelaces accepts standard genomic formats requiring alignment of ribosome profiling reads (BAM) and corresponding gene/ transcript annotations (GTF or BED). Shoelaces then guides the user through three main steps: (1) read filtering, (2) footprint identification and (3) P-site determination.

**Figure 1:**
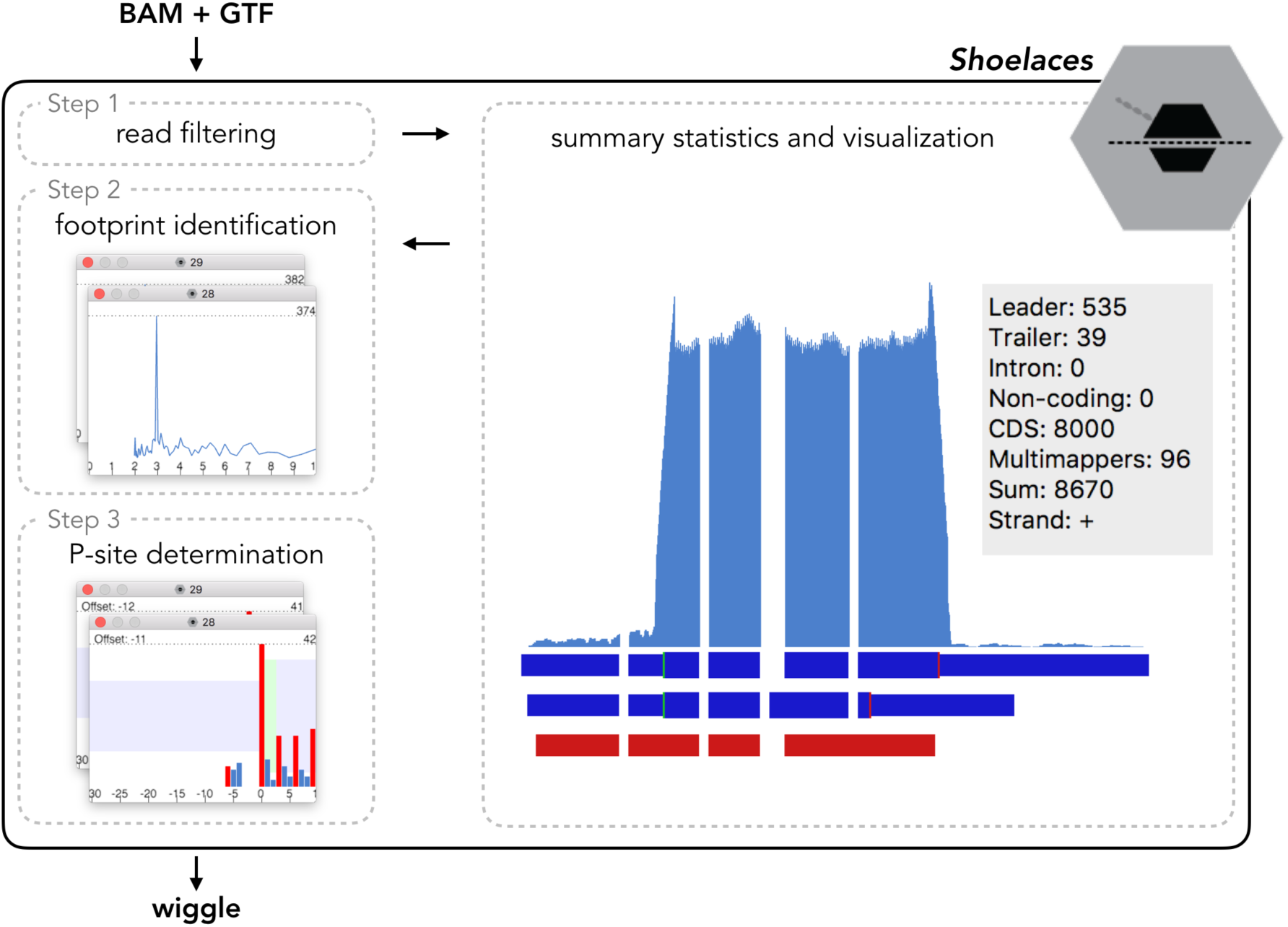
Shoelaces workflow. The toolkit accepts BAM and GTF files as input, filters reads, identifies translating lengths, determines P-site offsets and outputs tracks into wiggle format. Visual representation and summary statistics aid the processing steps.

In the initial step Shoelaces filters reads from noise regions. Here, users can optionally include an additional annotation file with regions (such as e.g. rRNA or repetitive elements) which will be masked from all further analyses. Specific genes can also be deselected during this step if certain outliers are undesired.

In the following step, Shoelaces automatically determines the correct footprint lengths. This is based on the intrinsic 3-nucleotide periodicity characteristic of ribosome-derived fragments as opposed to reads originating from other processes [7]. The periodicity is detected using discrete Fourier transform (see below) over the CDS regions of annotated genes. Lengths displaying periodicity are selected for further analysis. The rest is classified as noise but is available for further analysis by the user.

Finally, for each footprint Shoelaces determines the codon that is actively translated. A length dependent P-site offset is calibrated using change point analysis (see below) over the distribution of footprints surrounding start and stop codons of annotated genes. Based on this, Shoelaces will automatically suggest offsets and provide plots of the summed footprints over start and stop codons of all genes. In addition, ribosome footprints are known to map preferentially to the first nucleotide in the codon [1] and Shoelaces therefore displays the fraction of reads falling into each reading frame. Manual adjustment is also possible if deemed necessary by the user.

After confirming the selection of the suggested footprint lengths and offsets, the user can export the ribosome coverage into flat file format (wiggle) for further downstream analysis. Optionally, different footprint lengths can be exported into separate files. Separation by length can be useful for more specific analysis, such as e.g. detection of conformational changes of ribosome at certain positions [9, 10].

To aid the researcher, the GUI produces summary statistics and counts for individual genes and transcripts, as well as for the whole library. It provides an overview over how many reads of a given length fall into different genomic regions (CDSs, 5’ leaders, 3’ trailers and introns) as well as how many footprints are found over non-coding transcripts or mapping to multiple positions in the genome. Users can update the statistics after read length and offset selection to see how they change. Together, these give an indication of the quality of the library and how well the reads represent genuine ribosome protected fragments.

### Automatic selection of read lengths and offsets

An ideal-case scenario is presented in Fig. 2: the given footprint length is periodic (Fig. 2d), the metagene profiles have distinct peaks over start and before stop codons (Fig. 2a,b) and reads preferentially map to the first reading frame (Fig. 2c). However, library-specific biases can result in varying distributions of coding footprint lengths, as well as varying offsets (for various examples see Additional file 1). To take these biases into account, as well as to make processing large amounts of ribosome profiling data easy for the user, Shoelaces automatically suggests read lengths and offsets to be used.

**Figure 2:**
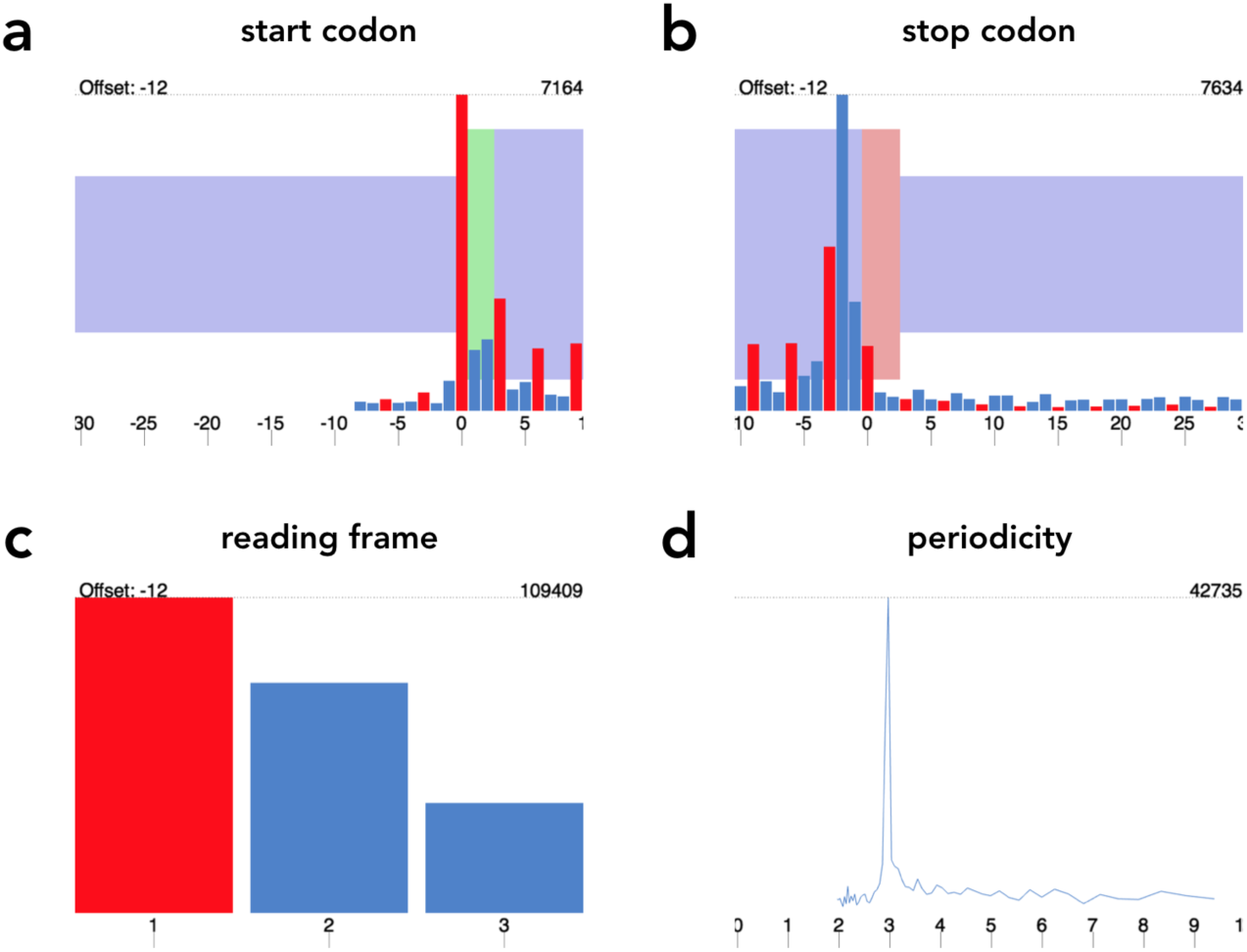
Read length and offset selection. In an ideal case scenario, the 3-nucleotide periodicity determines if the footprint length is coding (d), the peaks over start (a) and the last codon before stop (b) codons are used to calibrate offsets and most of the reads map to the first reading frame (c). Here, the plots demonstrate length 28 in human ribosome profiling sample (SRR493747, [15]). For more plots and datasets see Additional file 1.

### Selection of periodic lengths

For each fragment length, the 5’ ends of footprints mapping to the first 150 nucleotides of CDSs (by default from top 10% of protein-coding genes with highest coverage) are summed together. As the reads map preferentially to the first nucleotide of every codon, the periodic pattern will be conserved. The resulting vector is subject to discrete Fourier transform, and the fragment lengths whose highest amplitude corresponds to a period of 3 are considered to be periodic.

### P-site determination

For each fragment length, the distribution of 5’ ends of footprints surrounding start and stop codons (−30/+10 nucleotides) of protein-coding genes is calculated. The resulting window is subject to change point analysis, where for each adjacent position we calculate the difference of means. The maximum shift in means is assumed to correspond to the 5’ end of the footprints of initiating ribosomes and the distance from these to the P-site becomes the offset for that fragment length.

### Visualization

Shoelaces also allows for visual inspection of coverage over individual genes (or group of genes) of interest. Users can manually zoom in/out to adjust the view, inspect the summary statistics with and without using offsets, and export high quality figures and tracks for further analysis and visualization.

### Large-scale processing

For processing multiple libraries in bulk, a batch mode is available. For instance, for a number of same-batch libraries, one can be inspected visually, processing steps stored in an XML file and applied to the others. This additionally makes the processing easily reproducible later on. The processing can also be performed and fully automated from the command line allowing Shoelaces to be a part of a more comprehensive pipeline.

### Analysis of human ribosome profiling data

We analyzed 79 libraries of human ribosome profiling data from 12 studies [11, 12, 13, 14, 15, 16, 17, 18, 19, 20, 21, 22] and compared our read selection to the original, where applicable. Shoelaces retains up to 32% more data mapping to the coding regions of the genome (see Additional file 1, Table 1) than when originally processed. Non-periodic lengths are not selected, such as those that map primarily to 3’ trailers, suggesting that they might originate from e.g. mRNA-binding proteins, abundant in 3’ trailers, secondary structure or other sources of noise (Additional file 1, Figure 4).

## Conclusions

Shoelaces aims for an intuitive and streamlined processing of libraries from different studies and treatments, making them comparable and analysis easily reproducible. The precision in bringing the data to sub-codon resolution is especially important in studies on translational efficiency of different codons, but also allows for detection of translational events such as ribosomal pausing [23], stop codon readthrough [3] or frameshifting [6].

The automation and batch processing facilitate dealing with large amounts of data, while visualization features add to user-friendliness and allow for more specific analyses. As we demonstrate on human ribosome profiling data, Shoelaces retains more reads mapping to coding regions than arbitrary manual processing. Overall, Shoelaces is a comprehensive tool for ribosome profiling data processing, and should prove useful to anyone interested in small or large-scale studies on ribosome profiling.

### Availability of data and materials

The datasets analyzed in the current study are available in the Sequence Read Archive with accession numbers SRP038695 [11], SRP031501 [12], SRP002605 [13], SRP010679 [14], SRP012648 [15], SRP045257 [16], SRP014629 [17], SRP017263 [18], SRP053402 [19], SRP016143 [20], SRP029589 [21], SRP033369 [22].

The demo dataset is available together with the pipeline at https://bitbucket.org/valenlab/shoelaces.

### Funding

This work was supported by the Bergen Research Foundation and the Norwegian Research Council (#250049).

### Author’s contributions

Å B designed and implemented the software. KC implemented the algorithms for the method, tested the software, analyzed the data and wrote the manuscript. EV conceived the pipeline, guided the design and made critical revisions to the manuscript. All authors read and approved the final manuscript.

## Supplementary material

Additional file 1 — Analysis examples

Figure 1-3: Three different examples of offset selection (PDF file) for human ribosome profiling datasets: SRR493747 [15], treated with harringtonine and cyclohexamide; SRR1039861 [22], treated with cyclohexamide; SRR592961 [20], no drug. Table 1: Comparison of selected footprint lengths as originally in human ribosome profiling studies and Shoelaces. Figure 4: Comparison of reads mapping to different parts of transcript as selected by Shoelaces and the original manual selection (SRR493747 [15]).

